# A catalogue of Bilaterian-specific genes – their function and expression profiles in early development

**DOI:** 10.1101/041806

**Authors:** Andrea Krämer-Eis, Luca Ferretti, Philipp H. Schiffer, Peter Heger, Thomas Wiehe

**Affiliations:** Institut für Genetik, Universität zu Köln, 50674 Köln, Germany; Collège de France et Atelier de Bioinformatique, Universitè Pierre et Marie Curie, 75005 Paris, France

**Author notes:** Current address: The Pirbright Institute, Pirbright GU24 0NF, U.K. Current address: University College London, London WC1E 6BT, U.K.

## Abstract

Bilateria constitute a monophyletic group of organisms comprising about 99% of all living animals. Since their initial radiation about 540Mya they have evolved a plethora of traits and body forms allowing them to conquer almost any habitat on earth. There are only few truly uniting and shared morphological features retained across the phylum. Unsurprisingly, also the genetic toolkit of bilateria is highly diverged.

In the light of this divergence we investigated if a set of bilaterian-specific genes exists and, beyond this, if such genes are related with respect to function and expression patterns among organisms as distant as Drosophila, Caenorhabditis and Danio. Using a conservative pyramidal approach of orthology inference we collected a set of protein-coding genes which have orthologs in all major branches of Bilateria, but no homologs in non-bilaterian species. To characterize the proteins with respect to function, we employ a novel method for multi-species GO analysis and augmented it by a human-curated annotation based on an extensive literature search. Finally, we extracted characteristic developmental expression profiles for Bilateria from the extensive data available for three model organisms and we explored the relation between expression and function.

Among an initial set of several hundred candidates we identified 85 clusters of orthologous proteins which passed our filter criteria for bilaterian specificity. Although some of these proteins belong to common developmental processes, they cover a wide range of biological components, from transcription factors to metabolic enzymes. For instance, the clusters include myoD, an important regulator of mesodermal cell fate and muscle development, and prospero and several other factors involved in nervous system development. Our results reveal a so far unknown connection between morphological key innovations of bilateria, such as the mesoderm and a complex nervous system, and their genetic basis. Furthermore, we find typical expression profiles for these bilaterian specific genes, with the majority of them being highly expressed when the adult body plan is constructed. These observations are compatible with the idea that bilaterians are characterized by the unfolding of a new developmental phase, namely the transition of the larva to morphologically distinct adults.

**Author Summary:** Bilateria represent by far the largest and morphologically most diverse clade of all extant animals. The bilaterian radiation dates back to the so-called Cambrian explosion of species. Although bilateria show a large variety of very distinct body plans, they are also characterized by several common developmental and morphological traits, on which their monophyly is based. Here, we wanted to know whether these common phenotypic features may also have a shared and conserved genetic basis. To address this question we compared the proteomes of bilaterian and non-bilaterian species and extracted an initial set of a few hundred candidate proteins. Their underlying genes were further post-processed by means of orthology clustering, multi-species GO enrichment, expression analysis and extensive literature mining. This resulted in a thorough set of genes with roles in body morphology-, neuronal system‐ and muscle development, as well as in cell-cell signalling processes. This gene catalogue can be regarded as blue-print of a common bilaterian pheno‐ or morphotype and should contain highly interesting targets for further functional studies in model and non-model organisms.

## Introduction

Bilateria represent by far the largest monophyletic group in the animal kingdom. They comprise about 99% of the extant eumetazoans [1] and are classified into 32 phyla [2]. The taxon ‘Bilateria’ has been defined on the basis of morphological key innovations, namely bilateral symmetry, triploblasty, an enhanced nervous system and a complex set of cell types [3]. Probably the most striking observation is the existence of a large variety of body plans, accompanied by high morphological diversity. It is thought that the major bilaterian phyla emerged in a fast radiation during the early Cambrian, about 540 million years ago [1], but the appearance of the common bilaterian ancestor is controversial [4–6]. It has been suggested on the basis of conserved molecular patterning mechanisms, that the ‘ur-bilaterian’ was a complex vermiform animal with segmentation, two gut openings, lateral appendages, and coelomic space [4]. It is also assumed that the mesoderm and an anterior-posterior body axis were present in this animal [6] and thus are shared synapomorphies of the taxon. Still, it is difficult to derive unifying morphological properties for bilaterians by comparisons, as their descendants underwent extensive reduction and re-modelling of body forms in the lineages leading to extant crown clades. Nematodes, for example, lost their coelom, while insects lost some of the appendages found in crustaceans. On the other hand, insects gained ecological capacities by evolving the gills of their crustacean ancestors into wings [7]. Tetrapod limbs have evolved into the human hand, but also into fins during the evolution of cetaceans.

Despite these diversifications, comparisons at the molecular level reveal that many developmentally important genes are older than the ancestor of protostomes and deuterostomes. Among them are determinants of dorso-ventral and anterior-posterior patterning, eye formation, segmentation, and heart development [4,8–11]. In a recent study, Simakov et al (2013) estimated that up to 85% of the genes present in bilaterian crown clades were already in place in the molecular toolkit of the ur-bialterian. Deep comparative genomics, including non-bilaterian metazoans, revealed that the major developmental signaling pathways are already present in cnidarians, ctenophores and sponges [12–14], and must predate the origin of bilaterians. On the other hand, even highly conserved, and ‘important’, regulators can become lost, such as some Hox genes from *C. elegans* [15]. With roughly a billion years of divergence since the ur-bilaterian, chromosome fusions, fissions and re-arrangements, genome duplications, expansions and losses of gene families, as well as myriads of nucleotide substitutions have blurred the common molecular heritage. Given this situation, we wanted to compile a catalogue of those genes and proteins which have survived this mutational onslaught and can still be identified as orthologs across bilateria.

These proteins could allow us to glimpse at the molecular evolution in the bilaterian stem group, setting it apart from non-bilateria, and allow us to detect unifying molecular traits of the extant Bilateria [16]. To search for molecular traits that are present across divergent bilaterian species, we systematically compared their protein and coding gene complements with those of non-bilaterian species.

The genes which we could identify act in key developmental processes and show homologous expression profiles across comparable developmental stages in fish, fly and worm. Our findings lend support to the idea that changes in the evolution of development were key to the emergence of Bilateria.

## Results and Discussion

### Bilateria-specific clusters of orthologous proteins

The aim of our study was to identify genes that have newly emerged in the ‘ur-bilaterian’ and have been retained in its descendant species. Our rationale was that such genes constitute at least part of the genetic toolkit underlying common morphological or life-cycle features in bilateria. We restricted our search to those genes which most likely evolved in the bilaterian stem group, shown as a bold branch in the sketch of Fig 1a. Thus, they represent a novelty to the taxon or are highly divergent from non-bilaterian precursors. Our study is based on the comparison of ten bilaterian and seven non-bilaterian species with fully sequenced and annotated genomes. We downloaded 383, 586 protein sequences in total. About 70% (268, 252) are from bilateria, covering the three major clades Lophotrochozoa, Ecdysozoa and Deuterostomia, and about 30% (115, 334) are from non-bilateria (Table 1). We discarded all bilaterian sequences that had a reciprocal best BLAST hit below the threshold *E*-value of 10^−5^ with non-bilaterian sequences. We retained 13, 582 bilaterian specific candidate proteins which were grouped into 1, 867 clusters of orthologous proteins (’COPs’) using the OrthoMCL pipeline [17] (supp. Table S4).

**Figure 1.**
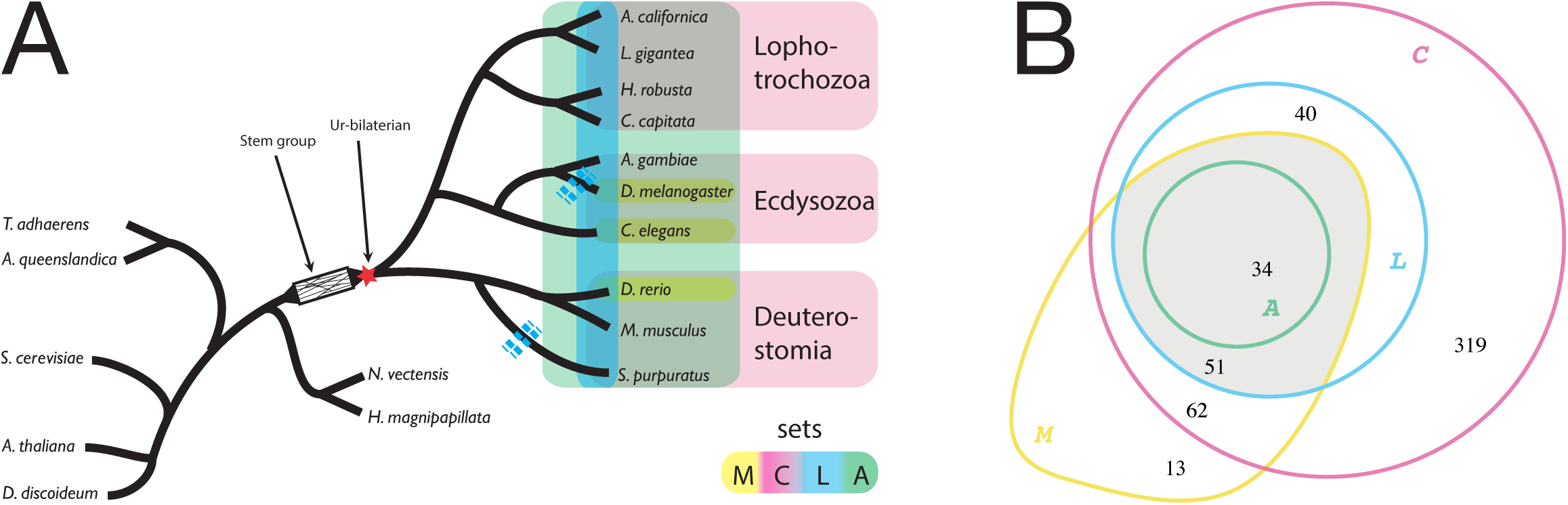
Panel A: Sketch of the genealogical relationship between the clades of Lophotrochozoa, Ecdysozoa and Deuterostomia and the species included in our analysis (Table 1). Legend: *M* – all <u>m</u>odel species are represented in a cluster; *C* – all <u>c</u>lades are represented; *L* – at most one loss event along the tree; *A* – <u>a</u>ll of nine bilaterian species are represented. Sketch of the ‘ur-bilaterian’ after [105]. Panel **B**: Absolute numbers of clusters of orthologous proteins (COPs) in the four sets. Gray region: detailed analyses are performed for 85 COPs in set *L*′ = *L* ⋂ *M*.

**Table 1.** Species considered to contrast bilaterian and non-bilaterian protein repertoires

| species | data source | no. proteins in database | no. proteins in all COPs | no. COPs w/ species<sup>1</sup> | no. proteins per COP<sup>2</sup> |
| --- | --- | --- | --- | --- | --- |
| **Bilateria** |  |  |  |  |  |
| Deuterostomia |  |  |  |  |  |
| *Danio rerio* | Ensembl | 41693 | 1190 | 428 | 2.78 |
| *Strongylocentrotus purpuratus* | NCBI | 42420 | 652 | 260 | 2.51 |
| *Mus musculus* | Ensembl | 40732 | 867 | 357 | 2.43 |
| Ecdysozoa |  |  |  |  |  |
| *Anopheles gambiae* | Ensembl | 13133 | 380 | 307 | 1.24 |
| *Drosophila melanogaster* | FlyBase | 23849 | 989 | 392 | 2.52 |
| *Caenorhabditis elegans* | WormBase | 25634 | 602 | 291 | 2.07 |
| Lophotrochozoa |  |  |  |  |  |
| *Aplysia californica* | NCBI | 1093 | 15 | 7 | 2.14 |
| *Capitella teleta* | JGI | 32415 | 642 | 376 | 1.71 |
| *Helobdella robusta* | JGI | 23432 | 420 | 325 | 1.29 |
| *Lottia gigantea* | JGI | 23851 | 492 | 337 | 1.46 |
| **Non-Bilateria** |  |  |  |  |  |
| *Amphimedon queenslandica* | NCBI | 9908 |  |  |  |
| *Arabidopsis thaliana* | TAIR | 28952 |  |  |  |
| *Dictyostelium discoideum* | Dictybase | 13426 |  |  |  |
| *Hydra magnipapillata* | NCBI | 17563 |  |  |  |
| *Nematostella vectensis* | JGI | 27273 |  |  |  |
| *Saccharomyces cerevisiae* | SGD | 6692 |  |  |  |
| *Trichoplax adhaerens* | JGI | 11520 |  |  |  |

1: number of COPs (out of $|C \cup M| = 519$ ), in which a given species is represented
2: average number of paralogs within a COP (number of proteins / number of COPs)

This strategy, based on reciprocal BLAST hits, has been used repeatedly to determine the evolutionary origin of genes and to assign a phylostratigraphic age [18–21]. One pitfall of this approach is the possibility to miss truly bilaterian-specific proteins with a conserved domain if proteins with a homologous domain are also present in non-bilaterians. For example, the C2H2 zinc finger protein CTCF has been described as bilaterian-specific [22]. Our search did not recover this protein, because multi-zinc finger proteins already exist in cnidarians and their conserved Cys/His residues and linker regions evoke a similarity above threshold. As a result, our approach will wrongly treat such proteins as older and not bilaterian-specific and, consequently, eliminate them. Similar errors may affect other proteins with extended conserved domains. However, proteins that pass the filter have a high probability to be true bilaterian novelties. On the other hand, genes may be wrongly assigned to phylostrata which are younger than bilateria. For instance, this could be a consequence of non-orthologous gene displacement (NOGD) [23] which can lead to the loss of genes in some bilaterian lineages. Considering both sources of error, it is therefore likely that we underestimate the number of truly bilaterian-specific proteins in our analysis.

We condensed the initial set of 1, 867 COPs by applying filters with different stringency for the taxa included (Fig 1) and obtained the following four sets

C: Each cluster must contain one or more representatives from each of the three major clades, Lophotrochozoa, Ecdysozoa and Deuterostomia. This set contains 506 clusters.
M: Each cluster must contain representatives from all model organisms, *D. rerio, D. melanogaster* and *C.elegans*. This set contains 160 clusters.
L: Each cluster contains representatives from all major clades, as in set *C*. However, only those clusters are permitted for which the representation of species is explained by at most one loss event along the species tree (set *C* does not have this restriction). This resulted in 125 clusters.
A: Each cluster contains representatives of all bilaterian species considered (no protein loss is admitted). This set has 34 clusters.

The bilaterian species are represented on average in 66.2% of all ortholog clusters in set *C* (average count of clusters among 506 total). In set *M*, this percentage is 80.9%, and in set *L* it is 91.2% (see Table 2), reflecting the levels of stringency of the filtering criteria.

**Table 2.**
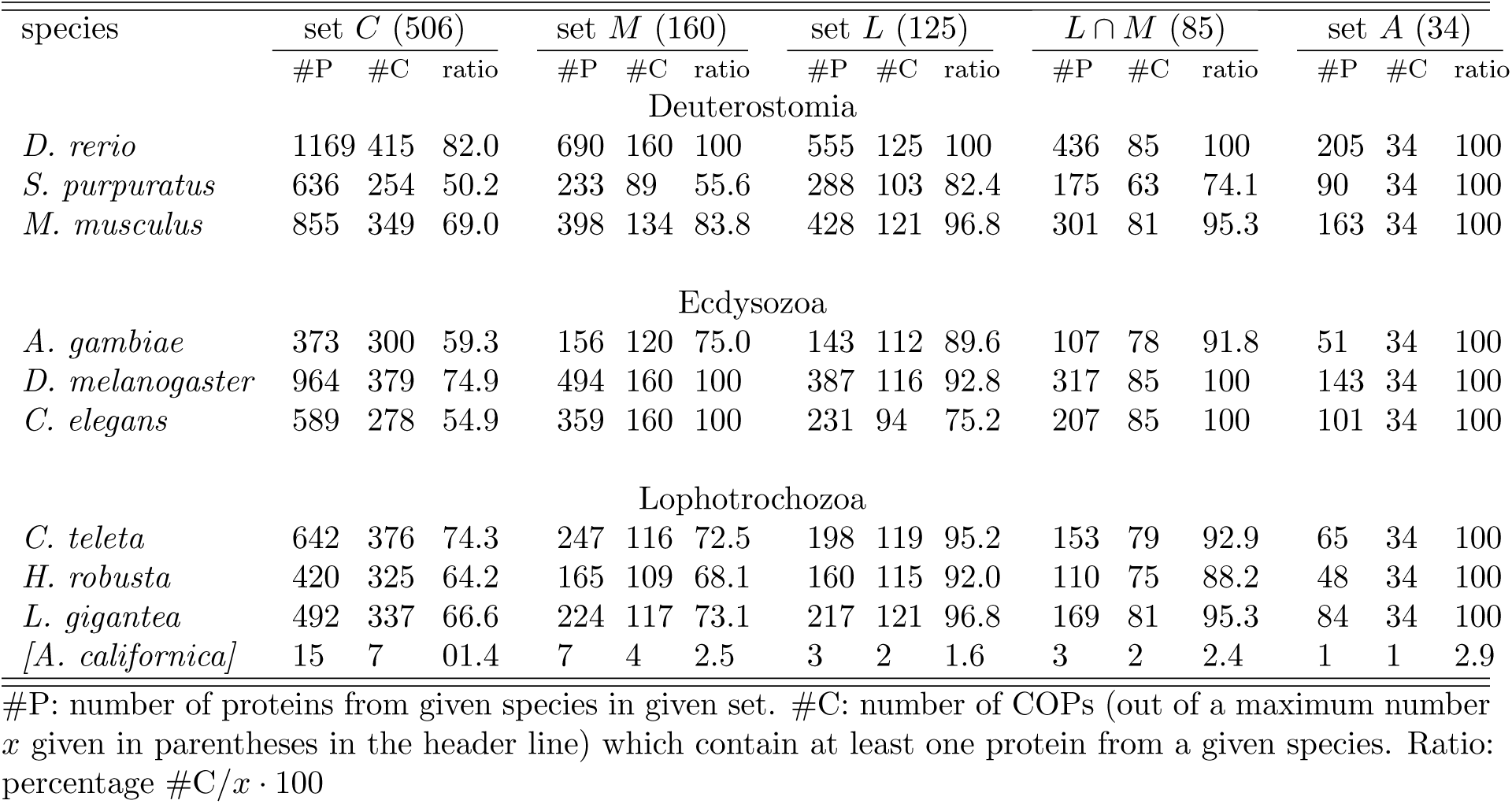
COP distributio;n in different data sets

A general drawback of our experimental procedure is its reliance on correctly identified and annotated genes. Since the majority of the organisms in our study are non-model organisms, they may suffer from incomplete or erroneous gene annotation. For example, the bilaterian species *A. californica* was omitted from downstream analyses because of its current low quality protein annotation. It is also possible that truly existing genes are missing from the non-bilaterian dataset due to annotation or prediction errors, leading to wrong inferences of bilaterian specificity. This error, however, is likely small because we interpret a pattern of presence and absence across several species, thereby reducing the impact of individual mistakes. Another limitation is the restriction to 16 genomes from which we infer bilaterian specificity. Although this is a very small fraction of the known metazoans, it contains all non-bilaterian genomes available at the time of the analysis and it constitutes a representative selection of bilaterian genomes. In particular, we included the model organisms *C. elegans, D. melanogaster*, and *D. rerio* in our analysis. These have very well curated and annotated genomes and are therefore helpful to find genes which are also functionally conserved since more than 500Myrs of independent evolution. To take advantage of the well curated annotation of the model organisms, we decided to use the intersection of sets *M* and *L* (termed *L*′ = *L* ⋂ *M*) for further analysis. Compared to set *L*, this intersection lacks 31 clusters missing a *C. elegans* ortholog and 9 clusters missing an ortholog from *D. melanogaster*, resulting in 85 final COPs. Disregarding *A. californica*, the least represented species in these clusters is *S. purpuratus* (genes of this species occur in 63 of 85 clusters). A fortiori, the model organisms (*C. elegans, D. melanogaster, D. rerio*) are represented in all clusters. On average, a (bilaterian) species is represented in 93.1% of the clusters. Non-bilaterian species which are not present in our initial dataset might possess true orthologs that we are not able to detect. To rule out this possibility, we verified by additional BLAST searches at NCBI that none of the 85 COPs had BLAST hits below an *E*-value of 10^−5^ in non-bilaterians or other eukaryotes (suppl Table S4). Again, non-orthologous gene displacement (NOGD) may obscure the fact that functional orthologs may escape detection when only working with sequence similarities.

As a result of our experimental design, but also due to different duplication histories, COPs contain differing numbers of paralogs from different species. On average we found most paralogs in Deuterostomia, followed by Ecdysozoa and Lophotrochozoa. The highest average paralog number is found for *D. rerio*, which might be a consequence of the additional genome duplication events in Teleosts [24]. Since the function of paralogs might differ from the last common ancestor, we extracted from each cluster in our set the most conserved ortholog for each species, according to the PhastCons conservation score [25], and discarded the less conserved paralogs. This reduction does not affect the number of clusters, but the number of proteins within a cluster. In our downstream analyses we considered both versions, the ‘most conserved orthologs’ (MCO) and the ‘all orthologs’ (AO) catalog for each cluster.

### Gene ontology and functional classification

#### GO terms reveal a link to development

To gain an overview over the biological functions of our set of proteins in set *L*′, we extracted and analyzed their associated GO terms. To quantify simultaneous enrichment in multiple species, we developed a novel generalized Fisher’s exact test for multiple species, which we term MSGEA (’Multi Species GO Enrichment Analysis’; Fig 2 and suppl. Fig S2). In contrast to standard GO enrichment analysis, MSGEA does not solely focus on the over-representation of GO terms in single species, but is able to detect GO terms enriched across several species, even if not over-represented in single species. Of special interest for our analysis were GO terms occurring across all three model organisms: such terms may indicate the conservation of biological function across large evolutionary time spans. To our knowledge, this novel method is the only test that is sensitive to coherent enrichment of GO terms across multiple species (see Methods). Applying MSGEA we extracted a list of GO terms which are likely associated to long-term retained functions of our candidate genes. Concentrating on the domain ‘biological process’ of the GO database, we find the terms ‘development’, ‘muscle’, ‘neuron’, ‘signaling’ and ‘regulation’ to be strongly over-represented (Fig. 3; suppl. Fig. S1; suppl. Table S6). This is compatible with the idea that key bilaterian innovations involved the central nervous system, muscles and complex cell types and tissues.

**Figure 2.**
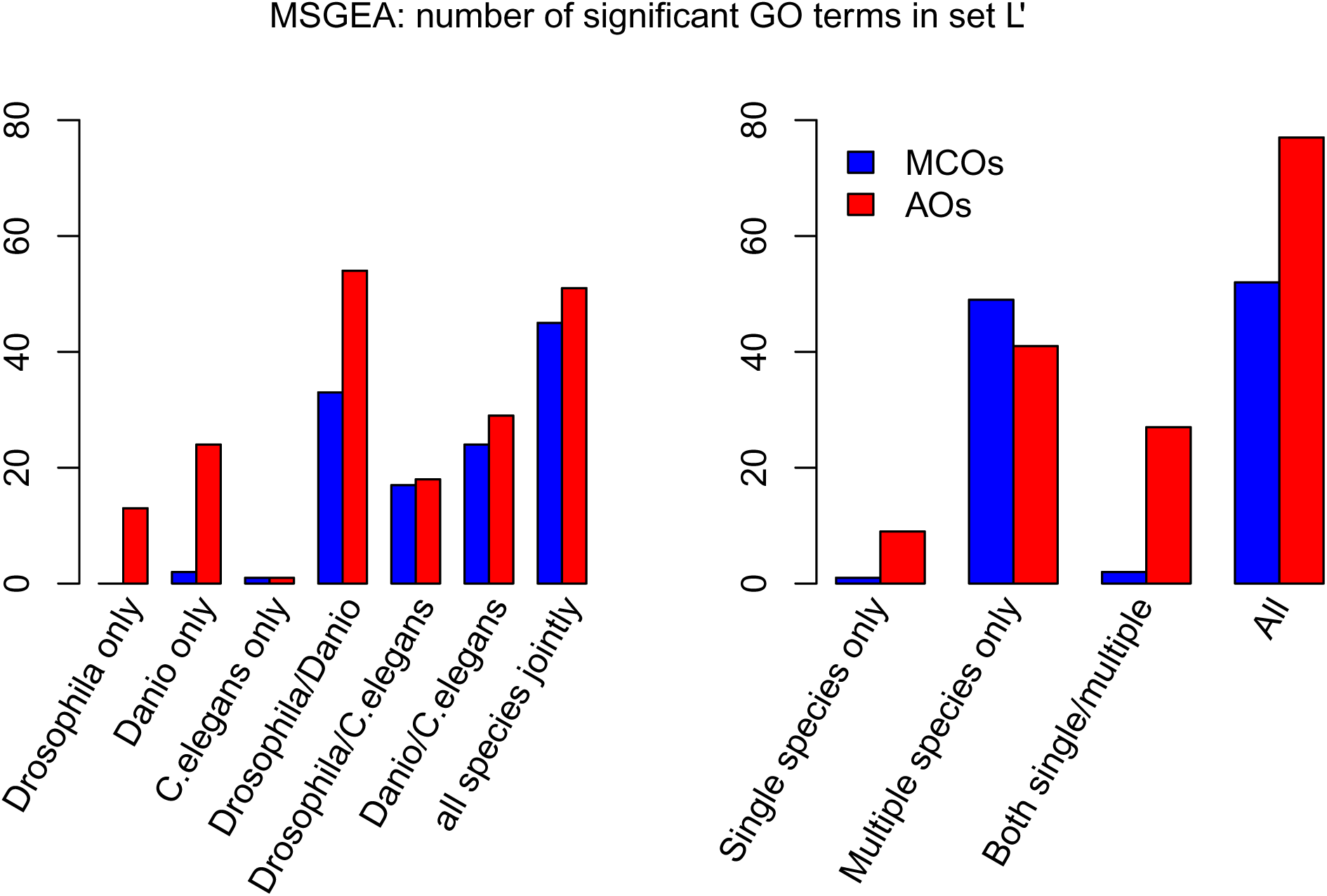
Number of significant GO terms identified with MSGEA in set *L*′ (*p* < 0.05). Blue histograms: considering all orthologs (AO) from a cluster; red histograms: only the most conserved orthologs (MCO) are considered in the MSGEA analysis. Left: number of significant terms found for each single‐ or multi-species enrichment analysis. Right: number of significant terms found only in single/multiple-species enrichment analyses and in both kind of analyses. The large amount of GO terms found only in multiple species analysis illustrates the power of MSGEA (see also suppl. Fig S2).

The terms identified through in silico analysis confirmed our hypothesis that prime candidates for bilaterian specific genes must be those which are acting during development.

#### Extraction of biological function through human curated literature mining

Since orthology is a strictly phylogenetic criterion, and since GO classification draws heavily on sequence homology, without necessarily reflecting conserved function, we decided to perform an in-depth human-curated literature search to collect additional evidence for the functional role of genes in COP clusters.

We mined the literature databases and compiled human curated functional descriptions, based on experimental evidence, for the proteins contained in set *L*′. We defined the six classes ‘Neuron related’, ‘Morphology related’, ‘Muscle related’, ‘Signaling related’, ‘Regulation related’ and ‘Others’ (suppl. Table S6). We used functional descriptions extracted from the literature to assign each of the 85 clusters to one of these six classes (Fig. 3). The manual extraction of species‐ and protein-specific functional information allowed us to compare and better interpret functions across the three model organisms.

**Figure 3.**
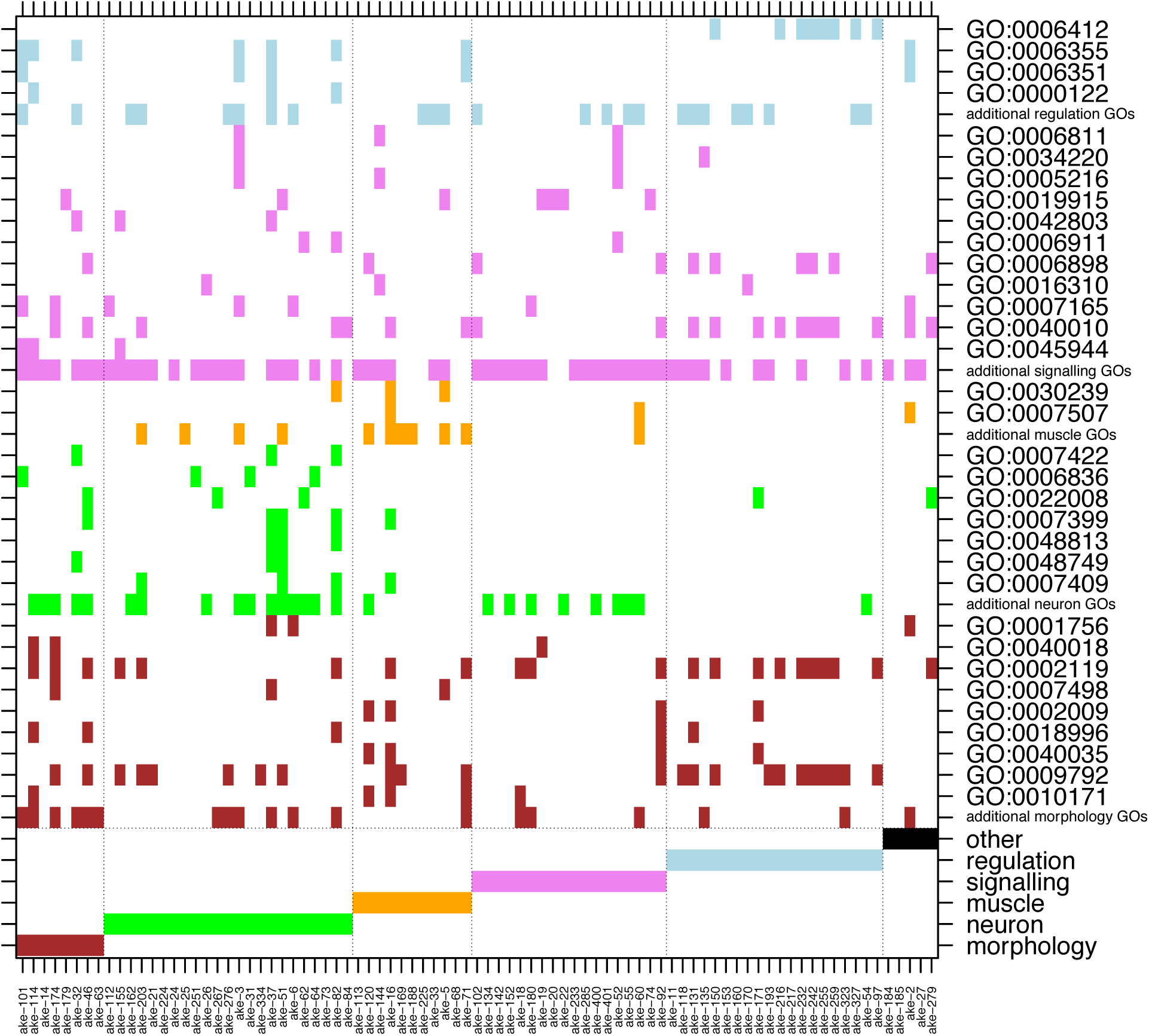
Significant GO terms identified in set *L*′ by applying ‘MSGEA’ (see text). The upper part of the figure shows the GO terms which occur in each COP (x-axis). Those terms which occur at least in three COPs are spelled out with their GO-ID, otherwise they are collected in the lines ‘additional … GOs’. Only non-generic GO terms of level 4 and higher are recorded. The bottom six rows show the assignment of COPs to functional classes based on human-curated literature mining. As can be seen from the color code, automatic GO and human-curated functional assignments do not always coincide.

To compare the human-curated classification with the one based on the GO database, we assigned all GO terms appearing in set *L*′ to one of the six classes above. Since a given protein may be associated with many GO terms there is no one-to-one relationship of the human curated annotation and the GO terms. For example, while the human curation assigns eight of the 85 clusters (the left-most columns in Fig 3) to the category ‘Morphology related’, morphology related GO terms occur in 33 clusters, scattered throughout the six classes. Similarly, signaling related GO terms are found in almost every cluster, while the class ‘Signaling related’ contains only 18 clusters. However, our manual curation is still broadly consistent with GO annotations. For example, GO terms associated to ‘muscle’ do occur in the class ‘Muscle related’ and GO terms related to ‘neuron’ are mostly found in the ‘Neuron related’ clusters.

#### ’Muscle-related’ COPs

One of our bilaterian-specific proteins is the basic helix-loop-helix (bHLH) transcription factor MyoD. MyoD and its paralogs Myf5, myogenin and MRF4 are well known as master regulators for muscle cell specification and differentiation in vertebrates (for a review see [26]). As so-called myogenic regulatory factors (MRFs), they coordinate the activities of many signaling proteins, chromatin modifiers and co-activators to control gene expression during myogenesis [26]. They are essential for the commitment of multi-potent somite cells to the myogenic lineage [27]. As in vertebrates, invertebrate MyoD orthologs are expressed in mesodermal tissue during early embryogenesis and are important for the specification of somatic muscle from these tissues [28,29]. Its functional conservation in different bilaterian phyla suggests that MyoD regulates muscle specification across bilateria. However, striated muscle is also present in non-bilaterian species, for example in the bell of cnidarian medusae or in fast-contracting tentacles of ctenophores [30,31]. On the basis of sequence and expression features, the bHLH protein JellyD1 has been proposed to regulate myogenesis and to be a functional MyoD ortholog in the hydrozoan jellyfish *Podocoryne carnea* [32]. However, more recent genome-wide surveys of bHLH transcription factors failed to detect MyoD orthologs in three other cnidarian species, questioning the general validity of the previous finding [33,34]. These observations and the uncertain classification of JellyD1 as MyoD ortholog [32] indicate that MyoD might be absent in cnidarians. Although supported by our data, additional work is necessary to confirm this conclusion. Taken together, the invention of MyoD in bilaterians might have facilitated the formation of elaborate gene circuits for muscle differentiation in these animals that are absent from non-bilaterians.

The functional contractile unit of striated muscle is the sarcomere. Z-disks on both sides delimit the sarcomere and serve as anchor for the evolutionarily old actin and myosin filaments. Several accessory proteins are required for proper sarcomere function. The giant protein titin, for example, functions as a myosin crosslinker and connects thick filaments axially to the Z-disk [35], the troponin complex (troponin T, I and C) tethers tropomyosin to actin in the absence of Ca2+, tropomyosin in turn is coiled around the actin helix and prevents its interaction with myosin if Ca2+ is absent (reviewed in [36]), and α-actinin, zormin and kirre tether the sliding filaments to the Z-disk (for review see [37]). Steinmetz et al. [38] concluded from sequence and domain arrangement comparisons that several sarcomere components are of bilaterian origin, the troponins C, I and T, titin, zormin and kirre. While a bilaterian origin of troponins is also a result of our study, we cannot reproduce the suggested origin for titin, zormin and kirre. All three proteins contain multiple immunoglobulin(-like) domains which are present also in non-bilaterian organisms, from sponges to cnidarians [39,40]. Similar to the CTCF example, the high similarity of immunoglobulin-like domains between bilaterians and non-bilaterians forced the exclusion of these proteins from the bilaterian-specific candidate set and explains the disagreement of our data with those of Steinmetz et al. [38].

Tropomyosin and actomyosin filaments exist also in cnidarian striated muscle [41]. Although the involvement of Ca2+ in jellyfish muscle contraction has been reported [42], their lack of Ca2+-sensing troponins indicates troponin-independent muscle contraction in these animals. The equivalent Ca2+ sensor of cnidarian muscle contraction is yet to be identified. As Steinmetz et al. [38] already pointed out, these findings suggest that a basal contractile apparatus existed in the common ancestor of cnidarians and bilaterians and was modified in lineage-specific ways in cnidarians and bilaterians.

#### ’Neuron-related’ COPs

To construct a complex nervous system, a hallmark of bilaterian crown clades, neurons have to migrate and grow axons to desired regions, for example muscles or nerve cells. While extracellular factors provide the cues for axon guidance, an intracellular machinery interprets these stimuli and directs appropriate cell movements. Microtubules and microtubule-associated proteins play a central role in these processes (for a review see [43]). We recovered in our screen for bilaterian-specific proteins two microtubule-associated proteins. One of them is the MAP1-family of microtubule-associated proteins that has three members in mammals and a single member, futsch, in *Drosophila melanogaster* [44]. The vertebrate ortholog MAP1B is highly expressed in the developing nervous system [45]. It is located at the distal part of extending axons and in growth cones, dynamic extensions of cytoplasm searching for their proper synaptic target during axon outgrowth and pathfinding. MAP1B promotes the nucleation, polymerization and stabilization of microtubules and is therefore required for these processes [43]. Similar functions have been described for the *Drosophila* ortholog futsch [46–48]. In addition, recent work from *Drosophila* showed that futsch is part of a membrane-associated microtubule organizing complex that controls synaptic dimensions and stability and thus the transmission speed of information [49].

Non-bilaterian animals such as cnidarians (and ctenophores) possess a diffuse nerve net consisting of ectodermal sensory and effector cells and endodermal ganglion cells. Although extracellular factors for axon guidance (netrins) have been discovered in cnidarians [50], it is not known how these extracellular stimuli regulate microtubule stability inside the cell, eg. by the action of microtubule-associated proteins, and how this contributes to axon pathfinding. In contrast to other microtubule-associated proteins, like kinesins and dyneins, whose origin predates the metazoan ancestor [51–53], the evolutionary history of the MAP1 family of microtubule-associated proteins has not been investigated so far. Our results indicate that MAP1 evolved in the ancestor of bilaterian animals, suggesting that bilaterians might have increased their capacity to wire a complex nervous system by a more sophisticated control of microtubule stability.

The other bilaterian-specific microtubule-associated protein recovered in our screen is tau. Tau (τ) is an intrinsically disordered protein with four microtubule-binding repeats and an N-terminal region that can associate with the cell membrane [54]. It is expressed mainly in the central and peripheral nervous system and is named after its ability to induce tubule formation [55]. Tau’s main function is to modulate the stability of axonal microtubules by binding to them. Tau-mediated remodelling and reorganization of dynamic microtubules in the growth cone is required for growth cone progression and axonal elongation, as well as for the recognition of guidance cues [56]. Tau’s binding to microtubules is regulated by phosphorylation (for review see [57]). Upon phosphorylation, tau detaches from microtubules and accumulates in neuronal cell bodies [58]. Hyperphosphorylation of tau can lead to the formation of tau-filaments which are involved in the pathogenesis of Alzheimer’s disease and other tauopathies [59]. Due to its medical relevance, there has been large interest in tauopathies, but the evolutionary history of tau has not been analysed so far. We found in our screen that tau evolved in the ancestor of bilaterians. In concert with the above mentioned results for MAP1/futsch this may indicate that the advent of bilaterians is characterized by a more elaborate control of microtubule stability and dynamics, with advantages for axon outgrowth and nerve cell connectivity that may have aided the development of complex nervous systems. These advantages must go beyond the generation of large axons, as the existence of giant axons in cnidarians is known for a long time [60]. At the same time, the tau example shows that a loss of microtubule control can be associated with pathological changes.

The third protein of the neuron-related group is prospero. Prospero is an atypical homeodomain transcription factor with orthologs described in other arthropods, nematodes and deuterostomes [61]. In *Drosophila*, prospero is well known for its key role in neuroblast differentiation and neural cell fate specification [62]. *Drosophila* neuroblast stem cells are arranged in a segmentally repeated pattern and give rise to all neurons and glia of the nerve cord by dividing asymmetrically into another neuroblast and a ganglion mother cell [63,64]. While the daughter neuroblast remains mitotically active for repeated asymmetric divisions, the ganglion mother cell divides only once more, and its daughter cells differentiate into nerve and glia cells that lose mitotic potential. Prospero acts during these processes as a binary switch between self-renewal and differentiation by promoting differentiation and repressing neuroblast-specific genes and cell cycle genes [65,66].

The vertebrate ortholog Prox1 has essential roles during morphogenesis of several organs, eg. of the lymphatic system, liver, or heart [67–69]. Prox1’s role as lymphangiogenetic factor is particularly well established. After the budding of endothelial cells from embryonic veins, Prox1 is required for their subsequent differentiation into lymphatic endothelial cells and therefore for the formation of a lymphatic vascular system [70]. Prox1 achieves this by regulating the expression of membrane receptor proteins and of cell adhesion and extracellular matrix molecules [71]. A conceptually similar function in the extension of small unicellular tubes has been described for the *C. elegans* prospero ortholog, ceh-26, that regulates genes mediating lumen extension in the tubules of the excretory canal cell [72]. Vertebrate Prox1 is further involved in the transition of neural progenitor cells from self-renewal to neuronal differentiation via direct suppression of Notch activity [73], a function remarkably similar to the processes in *Drosophila* nervous system development. Its expression in the nervous system of vertebrates, *Drosophila* and *C. elegans* and its importance in neurogenesis in *Drosophila* and vertebrates suggest that Prospero’s original function involved the transcriptional control of nervous system development and specification.

Besides prospero, a large number of transcription factors act in *Drosophila* and vertebrate neural development. These factors belong to different classes and include for example the Pax, Otx, and NK homeodomain protein families, bHLH proteins of the Atonal and Twist type, nuclear receptors such as retinoic acid receptors, and C2H2 zinc finger proteins of the Gli and Snail type (see [74], and references therein). Many of these neurogenic factors are present in poriferans and cnidarians and thus predate the evolution of bilaterians [74]. However, additional factors evolved in the ancestor of bilaterians when various neurogenic gene families were subject to another round of gene amplification (eg. Pax3/7, Engrailed, SoxA, SoxD, Neurogenin, Eomes) [74].

Our screen could not recover the majority of these known bilaterian-specific neurogenic factors. As they often originated from duplication events, their domains are inevitably similar to the domains present in the respective pre-bilaterian founder gene. A similarity above threshold leads to the exclusion of such genes from the dataset according to our filtering rules. Prospero, on the other hand, contains a highly divergent atypical homeodomain attached to a prospero DNA-binding domain (Homeo-prospero domain, pfam accession PF05044), a combination that is found only in bilaterians http://pfam.xfam.org/family/PF05044#tabview=tab7. Because of its unusual structural features, prospero is different from non-bilaterian homeodomain proteins and could be retained in the dataset. This result is in agreement with earlier studies that failed to detect homeodomain proteins of the PROS (prospero) class in cnidarians and ctenophores [75,76].

During nervous system development of cnidarians, three types of neurons arise from multipotent interstitial stem cells in *Hydra* [77] or from endoderm/ectoderm in *Nematostella* [78]. In contrast, neurogenesis in bilaterians is characterized by cell lineage expansion and diversification, involving multiple factors and dedicated neural progenitor cells [79]. Our results suggest that the bilaterian-specific homeodomain transcription factor prospero contributed to the sequence of nerve cell specifications via progenitor cells that is characteristic for bilaterians, and thus to the boost of complexity accompanying nervous system development in these animals.

### Expression profiles

To analyze expression of genes in set *L*′ across developmental stages and to compare it between model organisms, we used publicly available data for *D. melanogaster, D. rerio* and *C. elegans.* We normalized expression values - separately for each species and for each developmental stage - by the genome-wide average, and then log-transformed them.

#### Pooled profiles

To obtain a global picture of the expression profiles of genes in set *L*′, we calculated mean and median profiles for each of the three model organisms, shown as orange lines in Figs 4 and S3, top row. To see how expression of bilaterian-specific genes compares to background expression, we generated two different background sets for each species. The first is a random-background, obtained by randomly selecting genes from the model organism genomes and recording their expression (distribution shown as white boxplots in Fig 4). The second background, which we call the ‘matched-function’ background (orange boxplots), is a random collection of genes with GO terms matched to those found in set *L*′. The background distributions were constructed independently for each stage and for each set, and they may differ in size and content.

**Figure 4.**
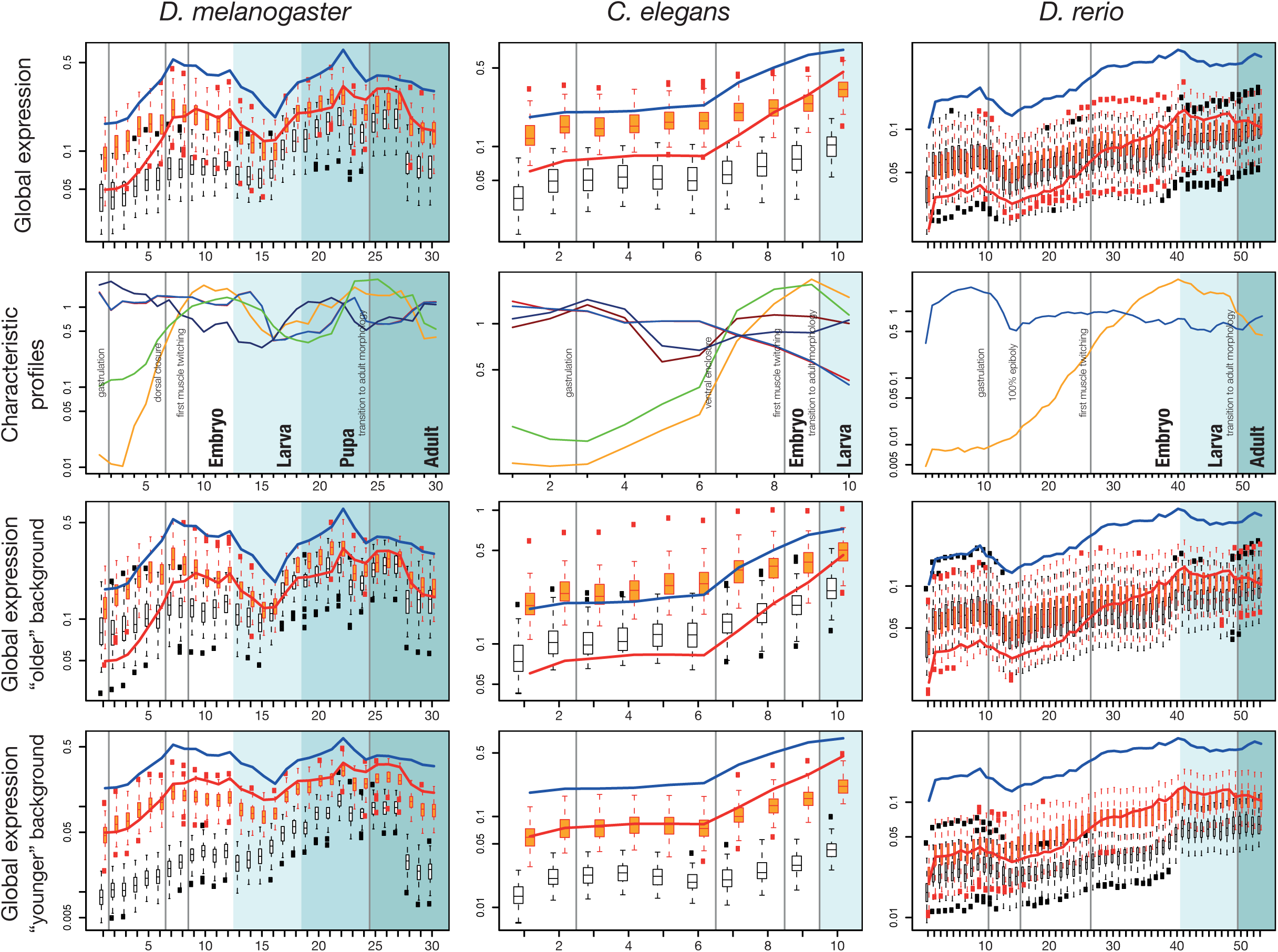
Expression profiles of different species. Top row: average (global) expression profiles for genes collected in set *L*′. Red line: pooled profiles for ‘all orthologs and paralogs’ (set ‘AO’). Blue line: profiles for the ‘most conserved orthologs’ (set ‘MCO’). The white and orange boxplots show the ‘random’ and ‘matched-function’ expression backgrounds, respectively (see text). Second row: Characteristic expression profiles. Different colors represent different clusters. Darker and lighter colors represent profiles which are similar among species. Bottom rows: distributions shown in boxplots are calculated from genes which are younger (phylostrata ≥ 7 according to the terminology of [19]) or older (phylostrata ≤ 7) than bilateria. Background shading (from white to dark grey) represents developmental stages embryo, larva, pupa (fly only) and adult. For a detailed explanation of the developmental stages see suppl. Tables S2 and S3.

We grouped the developmental stages from fertilized egg to the adult body into the rather coarse grained, yet comparable, phases: ‘Blastoderm’, ‘Gastrulation and Organogenesis’, ‘Hatching to Larva’ and ‘Adult’ (separated by vertical lines in Fig 4).

For all three species, the set *L*′ profiles roughly follow the shape of the background profiles across embryogenesis, however at a higher expression level (Fig 4). In *D. melanogaster*, expression of bilaterian-specific proteins is initially low, then rises until ‘dorsal closure’, after which it slighly decreases again. In *C. elegans*, expression is also initially low and then rises towards ‘ventral enclosure’. This pattern is mirrored in *D. rerio*, where expression peaks around the pharyngula stage. The profiles seen in *C. elegans* and *D. rerio* are congruent with the major cycle of tissue proliferation and differentiation during organogenesis in these species [80]. During ‘dorsal closure’ in *Drosophila*, ‘ventral closure’ in *C. elegans* and ‘pharyngula’ in *Danio*, the head, the nervous system and the bilaterally organized body develop [81]. In *Drosophila* we find a second expression peak during a second cycle of proliferation when the imago is formed in the pupa. This double-peaked profile seems to be characteristic for the fly, as observed previously [80,82]. The double peak pattern reflects two phases of increased cell proliferation during gastrulation and the transition from pupa to adult. The two peaks are also visible in the matched-function set, but they are less pronounced in the random-background set.

The matched-function profile largely resembles the profile of set *L*′. However, set *L*′ genes show distinctly low expression in the very early embryonic stages, a pattern which is consistently seen in all species. Another common feature is that expression of MCO genes is consistently higher than of AO genes (Fig 4). This can be explained by the fact that paralogs contained in set AO may be sub-functionalized, while the MCO genes are more likely to retain their original function.

#### Characteristic expression profiles

To further analyze the expression profiles, we clustered individual profiles based on their similarity and extracted a few typical ones for each species (Fig 4, bottom row). Methods exist to identify tissue-specific expression profiles, or to differentiate between profiles, but no standard techniques are available to detect enrichment of profiles across a set of genes. To overcome this limitation, we employed an ad hoc ‘profile enrichment’ strategy. Briefly, we considered an expression profile bilaterian-specific if similar profiles were significantly more abundant among bilaterian specific genes than among other genes (for details, see Methods). We extracted all statistically significant profiles and then clustered them to distill representative bilaterian-specific ones.

For each of the three species qualitatively similar patterns are encountered: an intersection of increasing expression profiles (yellow and light blue lines in Fig 4) with rather constant, or slightly decreasing profiles (dark red and dark blue lines). The intersection occurs right before the developmentally important events ‘dorsal closure’, ‘ventral enclosure’ and ‘pharyngula’. A second crossing of profiles can be seen in the fly, just before metamorphosis into the adult body.

#### Expression profiles stratified by function and age

We determined the mean expression profiles for the six functional classes ‐ morphology‐ (8 COPs), muscle‐ (11), neuron‐ (23), signaling‐ (18), regulation-related (20) and other genes (5) (suppl Fig S4). Since morphogenesis, neuron and muscle development are transitory processes we expect to find peaked expression patterns for the classes ‘Morphology related’, ‘Neuron related’ and ‘Muscle related’. Indeed, we find that the profiles of these three classes mirror the global trend of the pooled analysis, but with emphasized upward and downward peaks (Figure S5). In particular, expression before the onset of gastrulation is lower than for the genome-wide, matched-function backgound, a trend which is observed for all set *L*′ genes, but which is even more emphasized in this subclass of genes. On the contrary, the genes of the classes ‘Regulation related’ and ‘Signaling related’ display a rather constant profile, without the characteristic expression peaks of morphogenesis. This agrees with the interpretation that these genes conduct various, not necessarily transitory, functions.

For species as different as animals, plants and fungi, it has been repeatedly observed that timing and level of expression during development depend on the evolutionary age of a protein [19,21,81,83]. For instance, one recurrent finding was that phylostratigraphically older genes tend to show higher expression during early embryogenesis. To validate our proteins in this respect, we analysed whether the set *L*′ expression profiles might be more similar to the profiles of younger or of older genes and we calculated background distributions considering the age of genes. Genes which arose before Bilateria (phylostratigraphic level less than 7, according to the strata-numbers used by Domazet-Loso) were defined as ‘older’ and all others (level ≥ 7) as ‘younger’ [19]. Expression profiles of genes (AO) in set *L*′ match very well the profiles of the ‘younger’ genes with matched function (red lines and orange boxplots in Fig 4 and supp. Fig S3). All three species show quite low expression of young and of set *L*′ genes during early development. After ‘dorsal closure’ in *D. melanogaster*, ‘ventral closure’ in *C. elegans*, and ‘pharyngula’ in *D.rerio* expression increases and ‘older’ and ‘younger’ profiles converge.

The conserved genes in set *L*′ follow characteristic expression patterns during development, which are shared between species separated by a billion years of divergence. In particular, we observe a characteristic expression peak towards the end of larval development, indicating that this is also a genetically singular phase in development, unifying bilateria as distant as nematodes, insects and teleosts. The single (fish and nematode) and double peak (fly) patterns observed by us are in agreement with results from the modEncode project [80,82]. Starting from co-expression analyses, the authors arrived at a set of orthologous genes acting in comparable stages of *D. melanogaster* and *C. elegans* development. The simpler, single peaked pattern in *C. elegans* and *D. rerio* reflects up-regulation at ventral enclosure and pharyngula stages, respectively. Thus, the expression profiles in these organisms reflect their life-cycle, displaying shared peaks when the adult body-plan is assembled. This phase, dominated by morphogenesis, has been interpreted as the phylotypic stage in both organisms [84,85]. In the holometabolous fly the situation is more complex: a body morphology is built twice, for the larva and then for the adult. The two expression peaks we found for our proteins reflect this fact with upregulation during both phases of cell proliferation.

Neither the body form nor the developmental details of the ur-bilaterion are known with certainty. Davidson [86] suggested that it must have been an organism with ‘maximal indirect development’, with a primary larva which was morphologically very different from the adult [87]. It has also been claimed that stem-group bilaterians were minute planktotrophic creatures, similar to such primary larva and that the evolution of development into adult forms was a key transition for early bilateria [88,89]. However, based on studies of the morphology and development of acoel flatworms, it is now thought that the ur-bilaterion more likely resembled a planula-like organism with direct development [90]. The phylogenetic position of the Xenacoelomorpha and the enigmatic Xenoturbellids, an outgroup to other bilaterians, also suggest that mesoderm and establishment of three body axes were already characteristics for the ur-bilaterion [6]. The expression patterns of set *L*′ genes, peaking towards the end of development, could be a remnant of a phase in evolutionary history when a planula-like adult originated in the bilaterian stem group. In modern, highly derived crown-clades the ensuing diversification of body plans is then built on this last conserved phase. Thus, the regulatory processes underlying the morphogenic organization of an adult bauplan might have contributed to the evolution of modern animals [91]. Crucial genes and gene regulatory networks [92] (GRNs), orchestrating developmental mechanisms, are preserved today as the molecular backbone of bilateria [93].

Both our own and the modEncode analyses highlighted proteins acting in a variety of functions in development [80]. It has been hypothesised that differences in gene regulatory processes and the deployment of GRNs may have been more important for the evolution of bilateria than the conservation of single key genes [92]. However, non-bilateria have a similar complex distribution and abundance of gene regulatory elements and systems [94]. Without discrediting the idea of the importance of GRNs in shaping morphological innovations, our data indicate that the emergence of clade-specific genes does as well have a pivotal role in opening new areas in morpho-space. In particular, our results indicate that bilateria are unified by developmental processes, which are shaped by a set of conserved and clade-specific genes.

## Materials and Methods

Using a comparative genomic approach, we identified genes shared among bilateria. To represent the spectrum of the different bilaterian clades we have selected species from the three major clades - Deuterostomia, Ecdysozoa and Lophotrochozoa - concentrating on species with a fully sequenced genome (Table 1). We included more species from Lophotrochozoa than from the other two clades, since well-annotated complete genome sequences from Lophotrochozoa model-organisms were still under-represented in public databases at the beginning of our studies. To obtain a high contrast between bilateria and non-bilateria, we used seven non-bilaterian species, derived from a previous study [22], as an exclusion criterion for the further analysis. Proteomes and data from InterProScan [95] and the Gene Ontology consortium [96] were retrieved from the sources cited in Table 1.

### BLAST and Clustering

To find orthologs we set up a stringent analysis pipeline. First, we searched for homologs in bilaterians using the stand-alone BLAST version 2.2.25+. The downloaded protein sequences were grouped and again reciprocally BLAST-ed with a cut-off *E*-value of ≤ 10^−5^. We excluded all proteins for which a homolog was identified in at least one non-bilaterian. Second, we verified the potential bilaterian-specific proteins using two different ortholog finders, InParanoid and OrthoMCL [17,97]. Both were rated highly in benchmarking studies which analyzed the performance of orthology-prediction methods [98,99]. For the purpose of detecting orthologs of very diverged species, we chose OrthoMCL version 2.0.2 and the associated MCL version 11-335, as they were shown to be more robust [98,99] than similar programs. Third, we grouped the ortholog clusters into four sets (A, L, M, C; Fig 1), satisfying different additional stringency conditions. Due to the poor sequence quality, we treated the Lophotrochozoon *A. californica* differently: where available we included *A. californica* orthologs, but we did not require them to be present in set A (’all species’).

The above data sets are ordered by degree of confidence in the bilaterian specificity of the protein. At the same time they reflect the level of conservation of the proteins across different species. Seeking to be conservative regarding bilaterian specificity on the one hand (set A being most conservative), but also to analyze a dataset which is as comprehensive as possible, we focused our further analyses on the intersection of sets *L* and *M* (called *L*′; Fig 1). This set comprises 85 clusters.

For the three model species *D. rerio, D. melanogaster* and *C. elegans* we extracted the UCSC tracks of basewise PhastCons conservation scores [25] calculated across insects, teleosts and nematodes, respectively. We used these scores to rank all genes in a cluster according to their fraction of strongly conserved sites, i.e. sites with PhastCons score > 0.99. We selected the highest-ranking one from each cluster as the ‘Most Conserved Ortholog’ (MCO). These genes are likely functional orthologs, based on the idea that strong conservation reflects long-term evolutionary (and functional) constraint and that neo‐ or subfunctionalized paralogs tend to diverge faster. We used the fraction of strongly conserved sites, instead of the average conservation score, since we are interested in the degree of conservation in function, not in sequence. We were reasoning that functional conservation is related to high conservation of alleles at functional sites, while the remaining bases can evolve fast. However, the two ways of measuring similarity are highly correlated on a genome-wide scale across all coding exons (*r* = 0.91 for *D. melanogaster*, 0.84 for *D. rerio* and 0.71 for *C. elegans* (supp. Fig S6). We performed all subsequent analyses on both sets: the set of ‘All Orthologs and paralogs’ (AOs) and the set of MCOs.

### Multi-Species Gene Enrichment Analysis

To examine function of the genes in set *L*′ by *in silico* methods, we obtained their gene ontology (GO) terms for the three model species from the GO database [96].

To identify terms enriched in bilaterian specific genes compared to the genomic background, we employed two strategies: the conventional single-species enrichment analysis based on Fisher’s exact test and a novel method, termed multi-species gene enrichment analysis (MSGEA) (manuscript in preparation). Standard single-species Fisher’s exact tests for enrichment cannot capture a signal of moderate, but joint, enrichment across different species. Instead, with MSGEA we compute an exact *p*-value for a pooled set of species. It works as follows. First, for each GO term we counted the number of descendants in the GO tree. Then, we computed the *p*-values for the terms occurring in any species and corrected for multiple testing [100]. More specifically, let *n*_*go*, *s*_ be the count (i.e., number of occurrences) of the GO term *go* in species *s* contained in our set and let *N*_*go*, *s*_ be the count for the whole genome. Furthermore, let *n*_*s*_ = Σ_*go*_ *n*_*go*,*s*_ and *N*_*s*_ = Σ_*go*_ *N*_*go*, *s*_. Assume that all species under consideration diverged at the same time and evolved independently after that (i.e. they have a starlike phylogeny). The null hypothesis of the MSGEA test is that a given GO term was not over-represented in the common ancestor. Therefore, a significant *p*-value means that the genome of the ancestor was already enriched in the GO term considered. Being conservative, assume that the null distributions of *n*_*go*, *s*_ are independent hypergeometrics with parameters *N*_*go*, *s*_, *n*_*s*_ and *N*_*s*_ in each species. The enrichment statistics *X*_*go*_ is then the sum of the normalized enrichments of *n*_*go*, *s*_ across species, i.e. the sum of the z-scores:

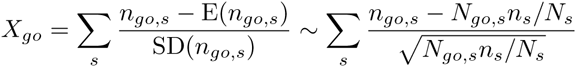

where a Poisson approximation is used to define the score. Finally, the *p*-value is the probability

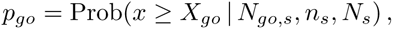

where the distribution of x follows from the hypergeometric distributions of the *n*_*go*, *s*_ with the above parameters.

For a single species, this test coincides with the standard one-tailed Fisher’s exact test for GO enrichment, and therefore is consistent. The exact estimation of *p*-values is computationally intensive; an optimized code, written in C, is available from the authors upon request.

Since MSGEA is based on the hypothesis of independent evolution of each lineage, it can be applied only to starlike phylogenies. For the situation considered here, this assumption is met, since the worm-fly-fish phylogeny is approximately starlike. We find that a considerable fraction (30-70%) of significant GO terms in our data set is detected by MSGEA, but not by single species enrichment.

### Detailed functional analysis of genes from model organisms

We employed biomaRt to mine literature databases for functional studies in model organisms with a focus on development: we used the Bioconductor module [101] biomaRt 2.19.3 to extract information for proteins present in cluster set L’. Wormbase release WS220 was queried for *C. elegans* proteins and ENSEMBL 75 for *D. melanogaster* and *D. rerio* proteins. We then searched extensively the literature for experimental validations of protein function. We grouped the results in six major categories, described in Results and Discussion and summarized in suppl. Table S6. These categories represent prominent molecular functions during embryogenesis and development. Based on the retrieved annotations we assigned proteins in set L’ to these categories.

### Expression

We retrieved expression data for different developmental stages in *D. melanogaster* [102,103], *D. rerio* [19] and *C. elegans* [104]. *D. rerio* data for adult stages were not used. The expression profiles were normalized by taking the logarithm to base 10 of the expression levels and then subtracting the log_10_ of the mean expression at each stage.

We devised the following test to find characteristic expression patterns for Bilateria. First, we computed Pearson’s correlation coefficient between expression profiles for different genes. For each profile in each set of bilateria-specific genes, we performed two types of tests: (i) a Mann-Whitney test on the distribution of correlations, comparing the correlation of the profile with other bilaterian genes versus the correlation with genome-wide profiles, in order to detect profiles that are more correlated with the ones of other bilaterian genes than with the rest of the genome; (ii) for each profile, we classified the remaining genes as “highly correlated” or “not highly correlated” in expression, using correlation thresholds of *r* = 0.5, 0.7 and 0.9; then, we tested for enrichment of correlated profiles among Bilateria by Fisher’s exact test.

A Benjamini-Hochberg correction for multiple testing was applied to the *p*-values resulting from these tests. All the significant profiles were clustered based on their correlation coefficients *r* by complete-linkage hierarchical clustering implemented in the R statistics software package, using 1 − *r* as distance measure and selecting clusters at height *h* = 0.75. After clustering, profiles were normalized by their average expression across stages, then averaged across each cluster. The resulting profile shapes are shown at the bottom of Figure 4.

### Expression versus function

We performed a randomisation test to explore if the mean (logarithmic) expression level of bilateria-specific genes could be explained by their function. For each developmental stage we checked if expression correlates with that of randomly selected genes possessing equal or similar GO terms, i.e. we checked if genes of different age, but similar ontology, would show similar expression profiles. We performed this analysis only on set L’.

For this purpose, we randomly sampled from the whole genome 100 sets with the same number of genes as are contained in set *L*′. We did this in two different ways. The first (“random-background”) was a random sampling of genes from the genome. After sampling, we computed the mean normalized expression of the selected genes for each stage. We repeated this procedure 100 times, and thus obtained a distribution of normalized means, represented as boxplots (in white) in Figs 4, S3, S5. The second was a “matched-function” sampling: first, for each GO term in the original set, we listed all genes with the same annotation and extracted from this list a number of random genes equal to the number of occurrences of the term; second, we pooled all the resulting lists; third, we extracted from this list a number of random genes equal to the number of genes in the original set. This way, we obtained sets of the same size and function (approximately, at least) as our original set.

We did this for the whole set L’ and for all subsets corresponding to the six functional categories described above (supp Fig S5).

### Age index of bilaterian-specific proteins

We used the phylostratigraphy from Domazet-Losšo et al. [19] for *D. melanogaster, D. rerio* and *C. elegans* to define two age-groups of proteins: (i) old proteins, with an origin in phylostrata 1 to 6; (ii) young proteins, with an origin in phylostrata 7 and higher. Proteins that we identified as bilateria-specific (i.e. the ones in our sets *A*, *L*, *M* and *C*) were excluded from the two groups, irrespective of their original phylostratigraphic classification. We repeated all the analyses described in the previous sections, and compared separately our bilateria-specific genes with the genome-wide background of older and of younger genes (Fig 4).

## Acknowledgments

This work was financially supported by a grant from the German Research Foundation (DFG-SFB680) to TW, and by a grant from the Volkswagen Foundation (Initiative for Evolutionary Biology) to PHS. We wish to thank Georgios Koutsovoulos and Sujai Kumar for advice with orthology inference and annotation.

---

Figure S1

Figure S2

Figure S3

Figure S4

Figure S5

Figure S6

Table S1

Table S2

Table S3

Table S4

Table S5

Table S6

## References

1. Marschall CR (2006) Explaining the Cambrian “explosion” of animals. Annual Review of Earth and Planetary Sciences 34: 355–384.

2. Nielsen C, Scharff N, Eibye-Jacobsen D (1996) Cladistic analyses of the animal kingdom. Biological Journal of the Linnean Society 57: 385–410.

3. Erwin D (2009) Early origin of the bilaterian developmental toolkit. Philos Trans R Soc Lond B Biol Sci 364: 2253–2261.

4. Hejnol A, Martindale M (2008) Acoel development indicates the independent evolution of the bilaterian mouth and anus. Nature 456: 382–386.

5. Philippe H, Brinkmann H, Copley R, Moroz L, Nakano H, et al. (2011) Acoelomorph flatworms are deuterostomes related to xenoturbella. Nature 470: 255–258.

6. Cannon J, Vellutini B, Smith J 3rd, Ronquist F, Jondelius U, et al. (2016) Xenacoelomorpha is the sister group to nephrozoa. Nature 530: 89–93.

7. Averof M, Cohen S (1997) Evolutionary origin of insect wings from ancestral gills. Nature 385: 627–630.

8. Saina M, Genikhovich G, Renfer E, Technau U (2009) BMPs and chordin regulate patterning of the directive axis in a sea anemone. Proc Natl Acad Sci U S A 106: 18592–18597.

9. Gauchat D, Mazet F, Berney C, Schummer M, Kreger S, et al. (2000) Evolution of ant p-class genes and differential expression of hydra hox/parahox genes in anterior patterning. Proc Natl Acad Sci U S A 97: 4493–4498.

10. Sinigaglia C, Busengdal H, Leclere L, Technau U, Rentzsch F (2013) The bilaterian head patterning gene six3/6 controls aboral domain development in a cnidarian. PLoS Biol 11: e1001488.

11. Sun H, Rodin A, Zhou Y, Dickinson D, Harper D, et al. (1997) Evolution of paired domains: isolation and sequencing of jellyfish and hydra pax genes related to pax-5 and pax-6. Proc Natl Acad Sci U S A 94: 5156–5161.

12. Moroz L, Kocot K, Citarella M, Dosung S, Norekian T, et al. (2014) The ctenophore genome and the evolutionary origins of neural systems. Nature 510: 109–114.

13. Srivastava M, Simakov O, Chapman J, Fahey B, Gauthier M, et al. (2010) The Amphimedon queenslandica genome and the evolution of animal complexity. Nature 466: 720–726.

14. Putnam N, Srivastava M, Hellsten U, Dirks B, Chapman J, et al. (2007) Sea anemone genome reveals ancestral eumetazoan gene repertoire and genomic organization. Science 317: 86–94.

15. Aboobaker A, Blaxter M (2003) Hox gene loss during dynamic evolution of the nematode cluster. Curr Biol 13: 37–40.

16. Dunn C, Leys S, Haddock S (2015) The hidden biology of sponges and ctenophores. Trends Ecol Evol 30: 282–291.

17. Li L, Stoeckert C Jr, Roos D (2003) OrthoMCL: identification of ortholog groups for eukaryotic genomes. Genome Res 13: 2178–2189.

18. Domazet-Loso T, Brajkovic J, Tautz D (2007) A phylostratigraphy approach to uncover the genomic history of major adaptations in metazoan lineages. Trends in Genetics 23: 533 - 539.

19. Domazet-Loso T, Tautz D (2010) A phylogenetically based transcriptome age index mirrors ontogenetic divergence patterns. Nature 468: 815–818.

20. Sestak M, Domazet-Loso T (2015) Phylostratigraphic profiles in zebrafish uncover chordate origins of the vertebrate brain. Mol Biol Evol 32: 299–312.

21. Cheng X, Hui J, Lee Y, Wan Law P, Kwan H (2015) A “developmental hourglass” in fungi. Mol Biol Evol 32: 1556–1566.

22. Heger P, Marin B, Bartkuhn M, Schierenberg E, Wiehe T (2012) The chromatin insulator CTCF and the emergence of metazoan diversity. Proc Natl Acad Sci USA 109: 17507–17512.

23. Koonin E (2003) Comparative genomics, minimal gene-sets and the last universal common ancestor. Nat Rev Microbiol 1: 127–136.

24. Taylor J, Braasch I, Frickey T, Meyer A, Van de Peer Y (2003) Genome duplication, a trait shared by 22000 species of ray-finned fish. Genome Res 13: 382–390.

25. Siepel A, Bejerano G, Pedersen J, Hinrichs A, Hou M, et al. (2005) Evolutionarily conserved elements in vertebrate, insect, worm, and yeast genomes. Genome Res 15: 1034–1050.

26. Berkes C, Tapscott S (2005) Myod and the transcriptional control of myogenesis. Semin Cell Dev Biol 16: 585–595.

27. Rudnicki M, Schnegelsberg P, Stead R, Braun T, Arnold H, et al. (1993) Myod or myf-5 is required for the formation of skeletal muscle. Cell 75: 1351–1359.

28. Michelson A, Abmayr S, Bate M, Arias A, Maniatis T (1990) Expression of a myod family member prefigures muscle pattern in Drosophila embryos. Genes Dev 4: 2086–2097.

29. Krause M, Fire A, Harrison S, Priess J, Weintraub H (1990) Cemyod accumulation defines the body wall muscle cell fate during c. elegans embryogenesis. Cell 63: 907–919.

30. Weber C, Alder H, Schmid V (1987) In vitro transdifferentiation of striated muscle to smooth muscle cells of a medusa. Cell Differ 20: 103–115.

31. Mackie GO, Mills CE, Singla CL (1998) Structure and function of the prehensile tentilla of euplokamis (ctenophora, cydippida). Zoomorphology 107: 319–337.

32. Mueller P, Seipel K, Yanze N, Reber-Mueller S, Streitwolf-Engel R, et al. (2003) Evolutionary aspects of developmentally regulated helix-loop-helix transcription factors in striated muscle of jellyfish. Developmental Biology 255: 216 - 229.

33. Simionato E, Ledent V, Richards G, Thomas-Chollier M, Kerner P, et al. (2007) Origin and diversification of the basic helix-loop-helix gene family in metazoans: insights from comparative genomics. BMC Evol Biol 7: 33.

34. Gyoja F, Kawashima T, Satoh N (2012) A genomewide survey of bHLH transcription factors in the coral Acropora digitifera identifies three novel orthologous families, pearl, amber, and peridot. Dev Genes Evol 222: 63–76.

35. Tskhovrebova L, Trinick J (2003) Titin: properties and family relationships. Nat Rev Mol Cell Biol 4: 679–689.

36. Brown J, Cohen C (2005) Regulation of muscle contraction by tropomyosin and troponin: how structure illuminates function. Adv Protein Chem 71: 121–159.

37. Luther P (2009) The vertebrate muscle z-disc: sarcomere anchor for structure and signalling. J Muscle Res Cell Motil 30: 171–185.

38. Steinmetz P, Kraus J, Larroux C, Hammel J, Amon-Hassenzahl A, et al. (2012) Independent evolution of striated muscles in cnidarians and bilaterians. Nature 487: 231–234.

39. Schaecke H, Rinkevich B, Gamulin V, Mueller IM, Mueller WEG (1994) Immunoglobulin-like domain is present in the extracellular part of the receptor tyrosine kinase from the marine sponge Geodia cydonium. Journal of Molecular Recognition 7: 273–276.

40. Rosa S, Powell A, Rosengarten R, Nicotra M, Moreno M, et al. (2010) Hydractinia allodeterminant alr1 resides in an immunoglobulin superfamily-like gene complex. Curr Biol 20: 1122–1127.

41. Groger H, Callaerts P, Gehring W, Schmid V (1999) Gene duplication and recruitment of a specific tropomyosin into striated muscle cells in the jellyfish Podocoryne carnea. J Exp Zool 285: 378–386.

42. Lin Y, Spencer A (2001) Localisation of intracellular calcium stores in the striated muscles of the jellyfish Polyorchis penicillatus: possible involvement in excitation-contraction coupling. J Exp Biol 204: 3727–3736.

43. Liu G, Dwyer T (2014) Microtubule dynamics in axon guidance. Neurosci Bull 30: 569–583.

44. Halpain S, Dehmelt L (2006) The MAP1 family of microtubule-associated proteins. Genome Biol 7: 224.

45. Tucker R, Garner C, Matus A (1989) In situ localization of microtubule-associated protein mRNA in the developing and adult rat brain. Neuron 2: 1245–1256.

46. Hummel T, Krukkert K, Roos J, Davis G, Klambt C (2000) Drosophila Futsch/22C10 is a MAP1B-like protein required for dendritic and axonal development. Neuron 26: 357–370.

47. Roos J, Hummel T, Ng N, Klambt C, Davis G (2000) Drosophila Futsch regulates synaptic microtubule organization and is necessary for synaptic growth. Neuron 26: 371–382.

48. Ruiz-Canada C, Ashley J, Moeckel-Cole S, Drier E, Yin J, et al. (2004) New synaptic bouton formation is disrupted by misregulation of microtubule stability in aPKC mutants. Neuron 42: 567–580.

49. Stephan R, Goellner B, Moreno E, Frank C, Hugenschmidt T, et al. (2015) Hierarchical microtubule organization controls axon caliber and transport and determines synaptic structure and stability. Dev Cell 33: 5–21.

50. Matus D, Pang K, Marlow H, Dunn C, Thomsen G, et al. (2006) Molecular evidence for deep evolutionary roots of bilaterality in animal development. Proc Natl Acad Sci USA 103: 11195–11200.

51. Wickstead B, Gull K (2006) A “holistic” kinesin phylogeny reveals new kinesin families and predicts protein functions. Mol Biol Cell 17: 1734–1743.

52. Wickstead B, Gull K (2007) Dyneins across eukaryotes: a comparative genomic analysis. Traffic 8: 1708–1721.

53. Wickstead B, Gull K, Richards T (2010) Patterns of kinesin evolution reveal a complex ancestral eukaryote with a multifunctional cytoskeleton. BMC Evol Biol 10: 110.

54. Morris M, Maeda S, Vossel K, Mucke L (2011) The many faces of tau. Neuron 70: 410–426.

55. Weingarten M, Lockwood A, Hwo S, Kirschner M (1975) A protein factor essential for microtubule assembly. Proc Natl Acad Sci U S A 72: 1858–1862.

56. Conde C, Caceres A (2009) Microtubule assembly, organization and dynamics in axons and dendrites. Nat Rev Neurosci 10: 319–332.

57. Ikezu S, Ikezu T (2014) Tau-tubulin kinase. Front Mol Neurosci 7: 33.

58. Lindwall G, Cole R (1984) Phosphorylation affects the ability of tau protein to promote microtubule assembly. J Biol Chem 259: 5301–5305.

59. Alonso A, Grundke-Iqbal I, Iqbal K (1996) Alzheimer’s disease hyperphosphorylated tau sequesters normal tau into tangles of filaments and disassembles microtubules. Nat Med 2: 783–787.

60. Mackie GO (1989) Evolution of cnidarian giant axons. In: Anderson PAV, editor, Evolution of the First Nervous Systems, Boston, MA: Springer. pp. 395–407. doi:10.1007/978-1-4899-0921-3_29. URL http://dx.doi.org/10.1007/978-1-4899-0921-3_29.

61. Elsir T, Smits A, Lindstrom M, Nister M (2012) Transcription factor PROX1: its role in development and cancer. Cancer Metastasis Rev 31: 793–805.

62. Doe C, Chu-LaGraff Q, Wright D, Scott M (1991) The prospero gene specifies cell fates in the Drosophila central nervous system. Cell 65: 451–464.

63. Broadus J, Skeath J, Spana E, Bossing T, Technau G, et al. (1995) New neuroblast markers and the origin of the aCC/pCC neurons in the Drosophila central nervous system. Mech Dev 53: 393–402.

64. Doe C, Technau G (1993) Identification and cell lineage of individual neural precursors in the Drosophila CNS. Trends Neurosci 16: 510–514.

65. Choksi S, Southall T, Bossing T, Edoff K, de Wit E, et al. (2006) Prospero acts as a binary switch between self-renewal and differentiation in Drosophila neural stem cells. Dev Cell 11: 775–789.

66. Li L, Vaessin H (2000) Pan-neural Prospero terminates cell proliferation during Drosophila neurogenesis. Genes Dev 14: 147–151.

67. Wigle J, Oliver G (1999) Prox1 function is required for the development of the murine lymphatic system. Cell 98: 769–778.

68. Sosa-Pineda B, Wigle J, Oliver G (2000) Hepatocyte migration during liver development requires prox1. Nat Genet 25: 254–255.

69. Risebro C, Searles R, Melville A, Ehler E, Jina N, et al. (2009) Prox1 maintains muscle structure and growth in the developing heart. Development 136: 495–505.

70. Wigle J, Harvey N, Detmar M, Lagutina I, Grosveld G, et al. (2002) An essential role for prox1 in the induction of the lymphatic endothelial cell phenotype. EMBO J 21: 1505–1513.

71. Hong Y, Harvey N, Noh Y, Schacht V, Hirakawa S, et al. (2002) Prox1 is a master control gene in the program specifying lymphatic endothelial cell fate. Dev Dyn 225: 351–357.

72. Kolotuev I, Hyenne V, Schwab Y, Rodriguez D, Labouesse M (2013) A pathway for unicellular tube extension depending on the lymphatic vessel determinant prox1 and on osmoregulation. Nat Cell Biol 15: 157–168.

73. Kaltezioti V, Kouroupi G, Oikonomaki M, Mantouvalou E, Stergiopoulos A, et al. (2010) Prox1 regulates the notch1-mediated inhibition of neurogenesis. PLoS Biol 8: e1000565.

74. Galliot B, Quiquand M (2011) A two-step process in the emergence of neurogenesis. Eur J Neurosci 34: 847–862.

75. Ryan J, Burton P, Mazza M, Kwong G, Mullikin J, et al. (2006) The cnidarian-bilaterian ancestor possessed at least 56 homeoboxes: evidence from the starlet sea anemone, Nematostella vectensis. Genome Biol 7: R64.

76. Ryan J, Pang K, NISC Comparative Sequencing Program, Mullikin J, Martindale M, et al. (2010) The homeodomain complement of the ctenophore Mnemiopsis leidyi suggests that Ctenophora and Porifera diverged prior to the ParaHoxozoa. Evodevo 1: 9.

77. David C (2012) Interstitial stem cells in hydra: multipotency and decision-making. Int J Dev Biol 56: 489–497.

78. Richards G, Rentzsch F (2014) Transgenic analysis of a soxb gene reveals neural progenitor cells in the cnidarian Nematostella vectensis. Development 141: 4681–4689.

79. Kohwi M, Doe C (2013) Temporal fate specification and neural progenitor competence during development. Nat Rev Neurosci 14: 823–838.

80. Gerstein M, Rozowsky J, Yan K, Wang D, Cheng C, et al. (2014) Comparative analysis of the transcriptome across distant species. Nature 512: 445–448.

81. Ninova M, Ronshaugen M, Griffiths-Jones S (2014) Conserved temporal patterns of microRNA expression in Drosophila support a developmental hourglass model. Genome Biol Evol 6: 2459–2467.

82. Li J, Huang H, Bickel P, Brenner S (2014) Comparison of D. melanogaster and C. elegans developmental stages, tissues, and cells by modENCODE RNA-seq data. Genome Res 24: 1086-1101.

83. Quint M, Drost H, Gabel A, Ullrich K, Bonn M, et al. (2012) A transcriptomic hourglass in plant embryogenesis. Nature 490: 98–101.

84. Kimmel C, Ballard W, Kimmel S, Ullmann B, Schilling T (1995) Stages of embryonic development of the zebrafish. Dev Dyn 203: 253–310.

85. Schiffer P, Kroiher M, Kraus C, Koutsovoulos G, Kumar S, et al. (2013) The genome of romanomermis culicivorax: revealing fundamental changes in the core developmental genetic toolkit in nematoda. BMC Genomics 14: 923.

86. Davidson E, Peterson K, Cameron R (1995) Origin of bilaterian body plans: evolution of developmental regulatory mechanisms. Science 270: 1319–1325.

87. Peterson K, Cameron R, Davidson E (1997) Set-aside cells in maximal indirect development: evolutionary and developmental significance. Bioessays 19: 623–631.

88. Peterson KJ, Davidson EH (2000) Regulatory evolution and the origin of the bilaterians. Proceedings Of The National Academy Of Sciences Of The United States Of America 97: 4430–4433.

89. Chen J, Oliveri P, Li C, Zhou G, Gao F, et al. (2000) Precambrian animal diversity: putative phosphatized embryos from the Doushantuo Formation of China. Proc Natl Acad Sci U S A 97: 4457–4462.

90. Hejnol A, Martindale MQ (2008) Acoel development supports a simple planula-like urbilaterian. Philosophical transactions of the Royal Society of London Series B, Biological sciences 363: 1493–1501.

91. Blackstone N, Ellison A (2000) Maximal indirect development, set-aside cells, and levels of selection. J Exp Zool 288: 99–104.

92. Davidson E, Erwin D (2006) Gene regulatory networks and the evolution of animal body plans. Science 311: 796–800.

93. Erwin D (2000) Life’s downs and ups. Nature 404: 129–130.

94. Schwaiger M, Schonauer A, Rendeiro A, Pribitzer C, Schauer A, et al. (2014) Evolutionary conservation of the eumetazoan gene regulatory landscape. Genome Res 24: 639–650.

95. Zdobnov E, Apweiler R (2001) Interproscan-an integration platform for the signature-recognition methods in interpro. Bioinformatics 17: 847–848.

96. Ashburner M, Ball C, Blake J, Botstein D, Butler H, et al. (2000) Gene ontology: tool for the unification of biology. The Gene Ontology Consortium. Nat Genet 25: 25–29.

97. Alexeyenko A, Tamas I, Liu G, Sonnhammer E (2006) Automatic clustering of orthologs and inparalogs shared by multiple proteomes. Bioinformatics 22: e9–15.

98. Shaye D, Greenwald I (2011) OrthoList: a compendium of C. elegans genes with human orthologs. PLoS One 6: e20085.

99. Hulsen T, Huynen M, de Vlieg J, Groenen P (2006) Benchmarking ortholog identification methods using functional genomics data. Genome Biol 7: R31.

100. Yoav Benjamini YH (1995) Controlling the false discovery rate: A practical and powerful approach to multiple testing. Journal of the Royal Statistical Society Series B (Methodological) 57: 289–300.

101. Guberman J, Ai J, Arnaiz O, Baran J, Blake A, et al. (2011) BioMart Central Portal: an open database network for the biological community. Database (Oxford) 2011: bar041.

102. Graveley B, Brooks A, Carlson J, Duff M, Landolin J, et al. (2011) The developmental transcriptome of Drosophila melanogaster. Nature 471: 473–479.

103. Cherbas L, Willingham A, Zhang D, Yang L, Zou Y, et al. (2011) The transcriptional diversity of 25 Drosophila cell lines. Genome Res 21: 301–314.

104. Levin M, Hashimshony T, Wagner F, Yanai I (2012) Developmental milestones punctuate gene expression in the Caenorhabditis embryo. Dev Cell 22: 1101–1108.

105. Telford M, Lockyer A, Cartwright-Finch C, Littlewood D (2003) Combined large and small subunit ribosomal RNA phylogenies support a basal position of the acoelomorph flatworms. Proc Biol Sci 270: 1077–1083.

